# In vitro analysis of catalase and superoxide dismutase mimetic properties of blue tattoo ink

**DOI:** 10.1101/2022.01.23.477399

**Authors:** Jan Homolak

**Affiliations:** Department of Pharmacology, University of Zagreb School of Medicine, Zagreb, Croatia; Croatian Institute for Brain Research, University of Zagreb School of Medicine, Zagreb, Croatia

**Keywords:** tattoo, tattoo ink, oxidative stress, catalase, superoxide dismutase

## Abstract

Tattoo inks are comprised of different combinations of bioactive chemicals with combined biological effects that are insufficiently explored. Tattoos have been associated with oxidative stress; however, a recent N-of-1 study suggested that blue tattoos may be associated with suppressed local skin oxidative stress. The present study aimed to explore the attributes of the blue tattoo ink (BTI) that may explain its possible effects on redox homeostasis, namely the catalase (CAT) and superoxide dismutase (SOD)-mimetic properties that have been reported for copper(II) phthalocyanine (CuPC) – the main BTI constituent. Intenze™ Persian blue (PB) BTI has been used in the experiment. CAT and SOD-mimetic properties of PB and its pigment-enriched fractions were analyzed using the carbonato-cobaltate (III) formation-derived H_2_O_2_ dissociation and 1,2,3-trihydroxybenzene autoxidation rate assays utilizing simple buffers and biochemical matrix of normal skin tissue as chemical reaction environments. CuPC-based tattoo ink PB and both its blue and white pigment-enriched fractions demonstrate CAT and SOD-mimetic properties *in vitro* with effect sizes demonstrating a substantial dependence on the biochemical environment. PB constituents act as inhibitors of CAT but potentiate its activity in the biochemical matrix of the skin. CuPC-based BTI can mimic antioxidant enzymes, however chemical constituents other than CuPC (e.g. the photoreactive TiO_2_) seem to be at least partially responsible for the BTI redox-modulating properties.

## Introduction

The art of tattooing dates back to the earliest stages of tribal communities and the oldest known tattoo belongs to the famous mummy *Ötzi the Iceman* (3000 BC)[1]. Throughout history, the prevalence of tattoos varied; however, in the last decades, the practice of tattooing spread throughout the Western world and became a mainstream form of body art. Recent estimates suggest that up to 25% of Europeans under the age of 20 and up to 38% of Americans under the age of 29 bear at least one tattoo [2]. Despite the omnipresence of tattooing, the biomedical effects of tattoos remain poorly explored, possibly because tattoo inks are comprised of different combinations of many bioactive chemicals with combined biological effects that are challenging to explore, let alone predict. In general, tattoo inks contain: i) organic (e.g. azo or polycyclic aromatic) or inorganic pigments (e.g. titanium dioxide (TiO_2_), barium sulfate (BaSO_4_), iron oxide, chromium oxide); ii) binders (e.g. polyethylene glycol, polyvinylpyrrolidone); iii) solvents (e.g. water, alcohol); and iv) additives (preservatives, surfactants)[2]. Additionally, impurities (e.g. nitrosamines or formaldehyde) may be present as well, and the ink injected into the body may contain nickel and chromium particles shredded from the needle during tattooing [3]. During the procedure of tattooing, all the constituents are delivered into the dermis where they become 100% systemically bioavailable due to direct contact with the blood and lymph. Although the kinetics of different tattoo ink constituents is still unknown, it is assumed that soluble components undergo rapid systemic distribution, while the insoluble pigments are mostly retained in the area of injection and in the draining lymph nodes where they may exert biological effects [2].

Although substantial efforts have been made to better understand the biological effects of tattoo inks utilizing *in vitro* and *in vivo* models, the results demonstrate that the observed effects are strongly dependent on the chemical constitution of the ink and the toxicological model utilized in the study. For example, Falconi et al. reported reduced viability and expression of the procollagen α1 type I in primary human fibroblasts incubated with *Biolip 27* but not *Strong black* [4]. Regensburger et al. reported substantial variability in the potency of 19 commercially available tattoo inks in respect to their inhibitory effects on mitochondrial activity in primary human dermal keratinocytes exposed to UVA radiation [5]. Arl et al. compared the effects of blue, green, red, and black tattoo ink on cell viability and the generation of reactive oxygen species (ROS) and reported that incubation with red and green tattoo inks induced the most pronounced toxic effects on the human keratinocyte cell line [6]. Perplexing *in vivo* results have also been reported. For example, in studies on tattoo ink carcinogenicity, mice tattooed with red tattoo ink and exposed to ultraviolet radiation developed tumors faster (214 vs 224 days) and showed increased tumor growth rate in comparison with sham-tattooed mice [7]. In contrast, black tattoo ink was protective against ultraviolet radiation-induced squamous cell carcinoma, delaying the tumor onset by approximately 50 days in tattooed mice [8]. Altogether, the biological effects of different tattoo inks seem to be too complex and specific to provide foundations for inductive reasoning on the effects of tattoo inks in general. To better understand the biomedical consequences of tattooing, a substantial effort should be made to i) elucidate the biological effects of individual chemicals present in different tattoo inks, and ii) explore the synergistic, additive, or antagonistic effects of chemical constituents of different tattoo inks in model systems that resemble those found *in vivo*.

The present aim was to explore the properties of blue tattoo ink that may explain the recently reported observation that a blue tattoo was able to suppress local skin oxidative stress [9]. Oxidative stress is a pathophysiological condition that ensues as a consequence of the inability of a system to maintain the balance between the electrophilic and the nucleophilic arm of redox homeostasis [10]. Considering redox homeostasis is critical for normal cellular functioning, it is no surprise that oxidative stress has been recognized as an important etiopathogenetic factor and a promising pharmacological target in pathophysiological conditions of the skin [11,12]. Tattoo inks have generally been associated with increased levels of oxidative stress (e.g. [5,6,13–15]); however, this has so far only been supported by indirect findings from *in vitro* experiments and there is currently no direct evidence for tattoo-induced oxidative stress in humans. In contrast, there is some evidence indicating that blue tattoo ink may be able to reduce oxidative stress. In an N-of-1 study, skin tattooed with blue tattoo ink demonstrated increased surface reductive capacity and the interstitial and intracellular fluid-enriched capillary blood from the tattoo had an increased content of protein sulfhydryls, reductive capacity, and catalase (CAT) activity, and reduced lipid peroxidation in comparison with the sample obtained from nontattooed skin [9]. Copper(II) phthalocyanine (CuPC), the main constituent of blue tattoo inks, can both reduce and prevent lipid peroxidation in homogenates of the mouse brain, kidney, and liver and exerts a substantial protective effect in the deoxyribose degradation assay [16]. Furthermore, it has been reported that CuPC can act as a dual functional mimetic of CAT and superoxide dismutase (SOD), two important antioxidant enzymes and that this property may be responsible for its lipid peroxidation-suppressing effects [17].

The present study aimed to explore whether: i) a blue CuPC-based tattoo ink can act as a CAT and SOD mimetic *in vitro*; ii) CAT and SOD mimetic properties of blue tattoo ink are primarily present in the CuPC-enriched ink fraction; iii) blue tattoo ink and its CuPC-enriched and residual fractions potentiate or inhibit the effects of CAT and SOD in the complex biochemical matrix of normal skin tissue (i.e. in the presence of endogenous CAT, SOD, and regulators of their activity).

## Materials and methods

### Sample preparation

Intenze™ Persian blue tattoo ink (PB) (Intenze, USA) was used in the experiment. The ingredients declared on the official material safety data sheet included: H_2_O (The European Community number (EC): 231-791-2), BaSO_4_ (EC: 231-784-4), TiO_2_ (EC: 215-280-1), CuPC (EC: 205-685-1), glycerine (EC: 200-289-5), isopropyl alcohol (EC: 200-661-7), *Hamamelis Virginiana* L. extract (EC: 283-637-9). Diluted PB samples were obtained by v/v dilution in pre-defined ratios in double-distilled H_2_O (ddH_2_O; 0.055 μS/cm). Dye fractionation was done by differential centrifugation. PB was first spun down for 30 minutes at a relative centrifugal force (RCF) of 12879 x g, and then the same process was repeated twice with both the supernatant (blue fraction) and the pellet (white fraction). The supernatant of the blue pigment-enriched fraction and the pellet of the white pigment-enriched fraction were used for subsequent analyses.

### UV-Vis spectrophotometry

UV-Vis spectra were obtained by scanning the absorbance in the wavelength range from 220 nm to 750 nm using The NanoDrop® ND-1000 (Thermo Fisher Scientific, USA).

### Catalase-like activity

CAT solution was prepared by dissolving 1 mg of lyophilized bovine liver CAT powder (Sigma Aldrich, USA) in 10 ml of phosphate-buffered saline (PBS) (pH 7.4). CAT activity was measured using the method first described by Hadwan [18] and adapted in [19]. Briefly, the samples were incubated with 50 μl of the substrate solution (2-10 mM H_2_O_2_ in PBS) and the reaction was stopped by adding 150 μl of the Co(NO_3_)_2_ stop solution (5 mL Co(NO_3_)_2_ x 6 H_2_O (0.2 g in 10 mL ddH_2_O) + 5 mL (NaPO_3_)_6_ (0.1 g in 10 mL ddH_2_O) added to 90 mL of NaHCO_3_ (9 g in 100 mL ddH_2_O)). The concentration of H_2_O_2_ was determined indirectly by measuring the absorbance of the carbonato-cobaltate (III) complex ([Co(CO_3_)_3_]Co) at 450 nm using the Infinite F200 PRO multimodal microplate reader (Tecan, Switzerland). Due to interference, a unique baseline model was established for each sample by simultaneous incubation with substrate solutions of graded nominal concentrations between 1 and 10 mM H_2_O_2_ and the Co(NO_3_)_2_ stop solution. The amount of residual H_2_O_2_ was estimated from the model for each sample and each time-point. CAT activity was assessed indirectly based on permutation-derived estimates of the baseline values (t = 0 s) and final values (t_1_ = 60 s or 300 s)[20].

### Superoxide dismutase-like activity

The SOD-like activity was measured by assessing the inhibition of 1,2,3-trihydroxybenzene (THB) autoxidation rate [21,22]. Briefly, 5 μl of each sample was placed in a 96 well-plate and incubated with freshly pre-mixed THB working solution (64 μl of 60 mM THB dissolved in 1 mM HCl mixed with 3400 μl of 0.05 M Tris-HCl and 1 mM Na_2_EDTA adjusted to pH 8.2). THB autoxidation was measured by assessing the absorbance increment at 450 nm with repeated measurements obtained by the Infinite F200 PRO multimodal microplate reader (Tecan, Switzerland).

### Preparation of the skin tissue constituents as the reaction matrix

To test whether the effects observed *in vitro* would be affected by the presence of skin tissue constituents present *in vivo*, a rat skin homogenate was prepared. Briefly, a piece of skin from a single rat euthanized in deep anesthesia (70 mg/kg ketamine; 7 mg/kg xylazine) was dissected and stored at -80 °C. The animal was in the control (untreated) group of another experiment and the tissue was dissected after decapitation in concordance with the 3Rs concept [23] in order not to interfere with the experimental protocol. The animal study from which the tissue was obtained was approved by the Ethics Committee of The University of Zagreb School of Medicine (380-59-10106-18-111/173) and the Croatian Ministry of Agriculture (EP 186/2018). The skin was rapidly dissected from the surrounding adnexa and placed in 1000 μl of lysis buffer (150 mM NaCl, 50 mM Tris-HCl, 1 mM EDTA, 1% Triton X-100, 1% sodium deoxycholate, 0.1% SDS, 1 mM PMSF, protease inhibitor cocktail (Sigma-Aldrich, USA) and PhosSTOP phosphatase inhibitor (Roche, Switzerland) adjusted to pH 7.5) on ice. The tissue was homogenized using Microson Ultrasonic Cell Disruptor (Misonix, SAD), centrifuged for 10 min at 4 °C, and RCF of 12 879 × g and the supernatant was stored at −80 °C. For the acute experiments, 5 μl of the tissue homogenate was used per well, and for the pretreatment experiments, 45 μl of the skin homogenate was incubated with either 5 μl of dd H_2_O, or 5 μl of the sample (1:10 PB, 1:10 blue, and 1:10 white fraction) for 180 min at 37°C.

### Data analysis

Data were analyzed in R (4.1.0). In the experiments where multiple substrate concentrations and multiple substrate exposure times were used, CAT-like activity was analyzed using linear regression with enzymatic activity (permutation-derived estimates) defined as the dependent variable and sample, substrate concentration, and time defined as independent variables. Model assumptions were checked by visual inspection of the residual and fitted value plots. Model outputs were reported as point estimates of least-square means with accompanying 95% confidence intervals. Comparisons of the samples from the model were reported as effect sizes (differences of estimated marginal means with accompanying 95% confidence intervals). Alpha was set at 5% and p-values were adjusted using the Tukey method.

## Results

### Persian blue tattoo ink acts as a weak catalase and superoxide dismutase mimetic *in vitro*

PB demonstrated dilution-dependent CAT-like activity *in vitro*, with the 1:10 dilution showing the ability to dissociate ∼0.9 mM of H_2_O_2_ in 300 s on average, in substrate concentrations ranging from 6 to 10 mM (**Fig 1A**). PB (1:10) dissociated 0.6 mM of H_2_O_2_ in the presence of 10 mM H_2_O_2_ and 1 mM when incubated with 8 and 6 mM, indicating lower efficacy at high substrate concentrations. The 1:100 dilution of PB demonstrated no CAT-like activity. The largest tested PB concentration (1:10) exerted SOD-like activity as well, by reducing the rate of THB autoxidation in the first 300 s of the assay. After 300 s, the maximum suppressive capacity of 1:10 PB was reached and PB potentiated autoxidation (**Fig 1B**). Lower PB concentrations (1:100; 1:1000; 1:10 000) showed no SOD-like activity.

**Fig 1.**
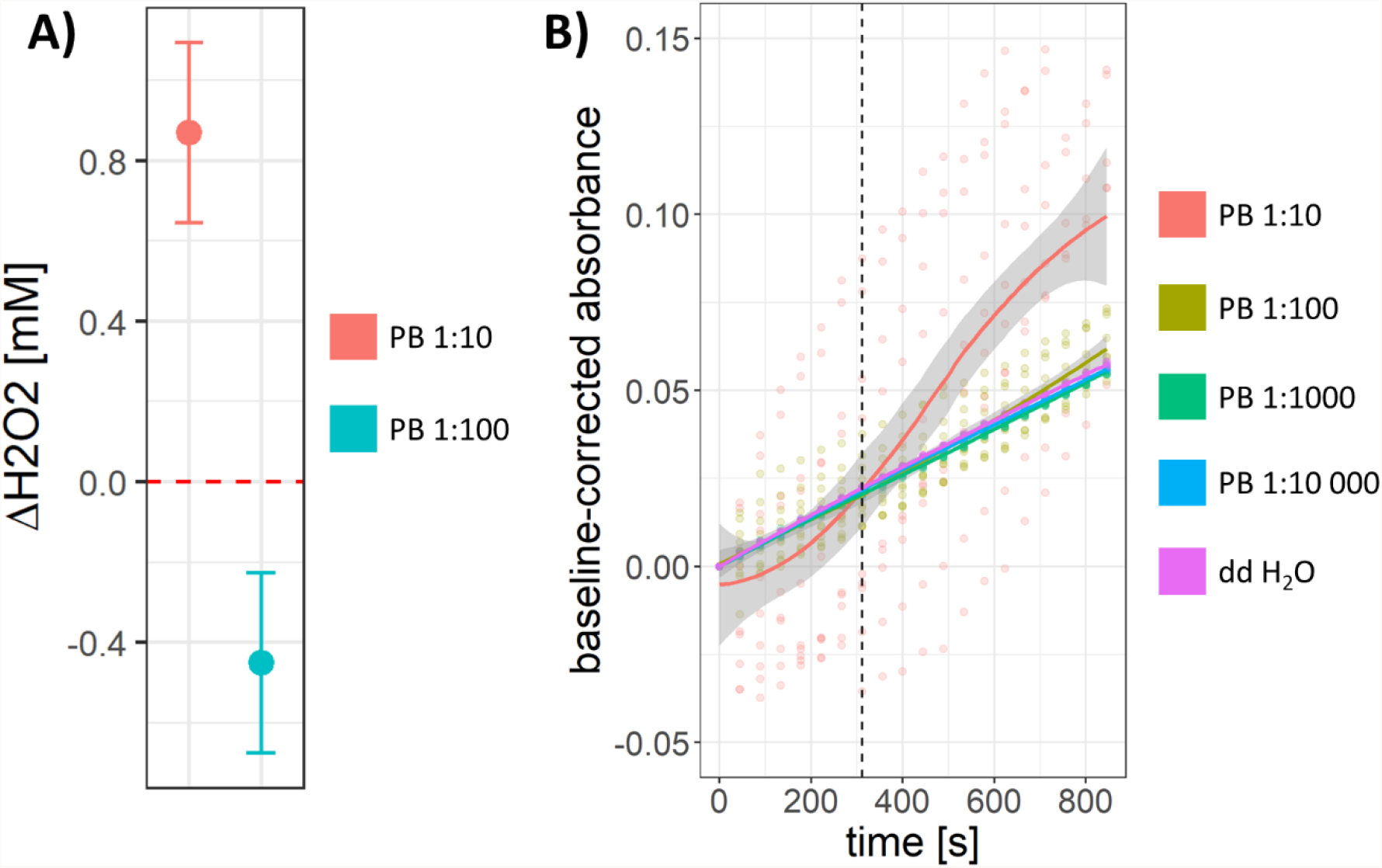
Catalase (CAT) and superoxide dismutase (SOD)-like activity of Persian Blue tattoo ink (PB) *in vitro*. A) Output of the model including the tested dilutions and substrate concentrations demonstrating dilution-dependent CAT-like activity of PB, with the 1:10 PB dilution acting as CAT mimetic (substrate concentrations: 6, 8, and 10 mM H_2_O_2_; t= 300 s). B) Dilution-dependent SOD-like activity of PB, with 1:10 PB demonstrating SOD-like properties at t < 300 s, and potentiation of 1,2,3-trihydroxybenzene autooxidation at t > 300 s. 1:100, 1:1000 and 1:10 000 dilutions show no SOD-mimetic activity.

### Blue and white fractions of the Persian blue tattoo ink show no catalase mimetic properties, but demonstrate divergent superoxide dismutase-like behavior

Centrifugation-based fractionation of PB yielded a blue CuPC-enriched fraction and a white fraction most likely enriched with TiO_2_ and BaSO_4_ (**Fig 2A**). UV-Vis spectra of fractionated samples suggested that CuPC was primarily present in the blue fraction, as evident by the presence of the Soret peak (B-band) and the Q-band with the Davydov splitting characteristic for the phthalocyanine derivatives [24,25] (**Fig 2B**). Neither the blue nor the white fraction demonstrated CAT-like activity *in vitro* (**Fig 2C-E**). Interestingly, "negative estimates of the activity” were obtained for the 1:10 dilution of both fractions indicating possible photocatalytic generation of H_2_O_2_ [26]. The observed effect was substrate concentration-dependent and more pronounced for the white fraction, which also demonstrated a pronounced time-dependence (**Fig 2C-E**). While the CuPC-enriched blue fraction showed no SOD-like activity at t < 300 s, the white fraction (1:10) potentiated THB autoxidation. At t > 300 s, the blue PB fraction (1:10) demonstrated SOD-like activity (**Fig 2F**).

**Fig 2.**
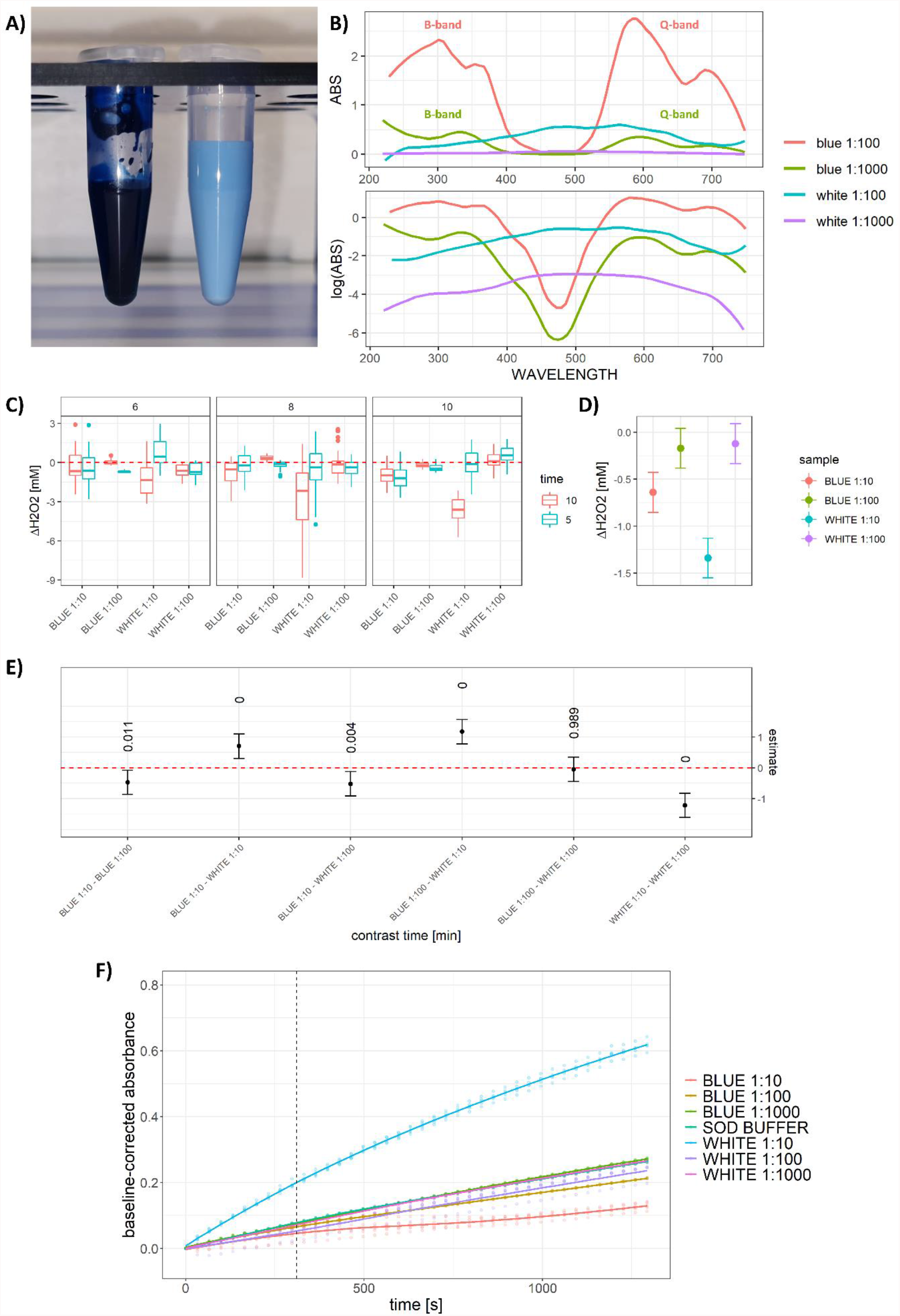
Catalase (CAT) and superoxide dismutase (SOD)-like activity of blue and white fractions of the Persian Blue tattoo ink (PB) *in vitro*. A) A representative image of the blue (left) and white (right) fractions of PB. B) Absorption spectra of the 1:100 (red) and 1:1000 (green) dilution of the blue fraction, and the 1:100 (turquoise) and 1:1000 (purple) dilution of the white fraction of PB. The absorption spectrum of the blue fractions demonstrates the Soret peak (B-band) and Q-band, strongly indicating that copper phthalocyanine was only present in the blue fraction of PB. C) Results demonstrating no CAT-like activity of different dilutions of either fraction *in vitro* (substrate concentrations: 6, 8, 10 mM H_2_O_2_; t_1_= 300 s; t_2_= 600 s). D) Output of the model including tested dilutions of the blue and white fraction, time, and substrate concentration, demonstrating no observed CAT-like activity. E) Comparison of the observed CAT-like activities of two dilutions of the blue and white fraction at different substrate concentrations. P-values are reported above estimates of differences of estimated marginal means accompanied by 95% confidence intervals. F) Dilution-dependent SOD-like activity of PB fractions. The 1:10 dilution of the white fraction potentiates autooxidation of 1,2,3-trihydroxybenzene at t < 300 s, while all other dilutions of both fractions show no pronounced effects. At t > 300 s, the 1:10 dilution of the blue fraction demonstrates pronounced, while the 1:100 dilution shows slightly less pronounced SOD-like activity. The 1:10 dilution of the white fraction shows a strong 1,2,3-trihydroxybenzene autooxidation potentiating effect.

### Persian blue tattoo ink and its blue and white pigment-enriched fractions inhibit catalase *in vitro* but potentiate its action in the presence of biochemical constituents of the skin

Apart from acting as CAT/SOD mimetics, PB constituents may modulate redox balance by affecting the activity of endogenous enzymes. In the presence of bovine liver CAT, the CuPC-enriched PB fraction acted as a weak CAT inhibitor with no evident dose-response, while PB and the white fraction exhibited a pronounced dose-dependent inhibition of the enzyme (**Fig 3A, B**). A different pattern was observed in the presence of biochemical constituents of the skin, where PB and its blue and white pigment-enriched fractions acted as potentiators of endogenous CAT, with the largest effect observed in the presence of the white fraction (**Fig 3C, D**). As the acute effects may not faithfully represent biochemical effects that may take place *in vivo*, an additional experiment was conducted in which the tested samples were first incubated with the biochemical constituents of the skin. Interestingly, following prolonged incubation (180 min at 37 °C), the pronounced effect of the white fraction was substantially attenuated, and the CuPC-enriched fraction induced the most pronounced effect on the activity of endogenous CAT (**Fig 3E, F**). In the presence of tissue constituents, both the blue and white fractions acted as SOD mimetics at t < 300 s, while there was no difference between the effect of PB and the control condition (**Fig 3G**). After the 300 s time-point, the CuPC-enriched fraction demonstrated stable SOD mimetic activity, while the white fraction potentiated THB autoxidation (**Fig 3G**). Following the prolonged incubation at 37 °C, SOD potentiating effects of PB and its blue and white fractions were completely lost (**Fig 3H**).

**Fig 3.**
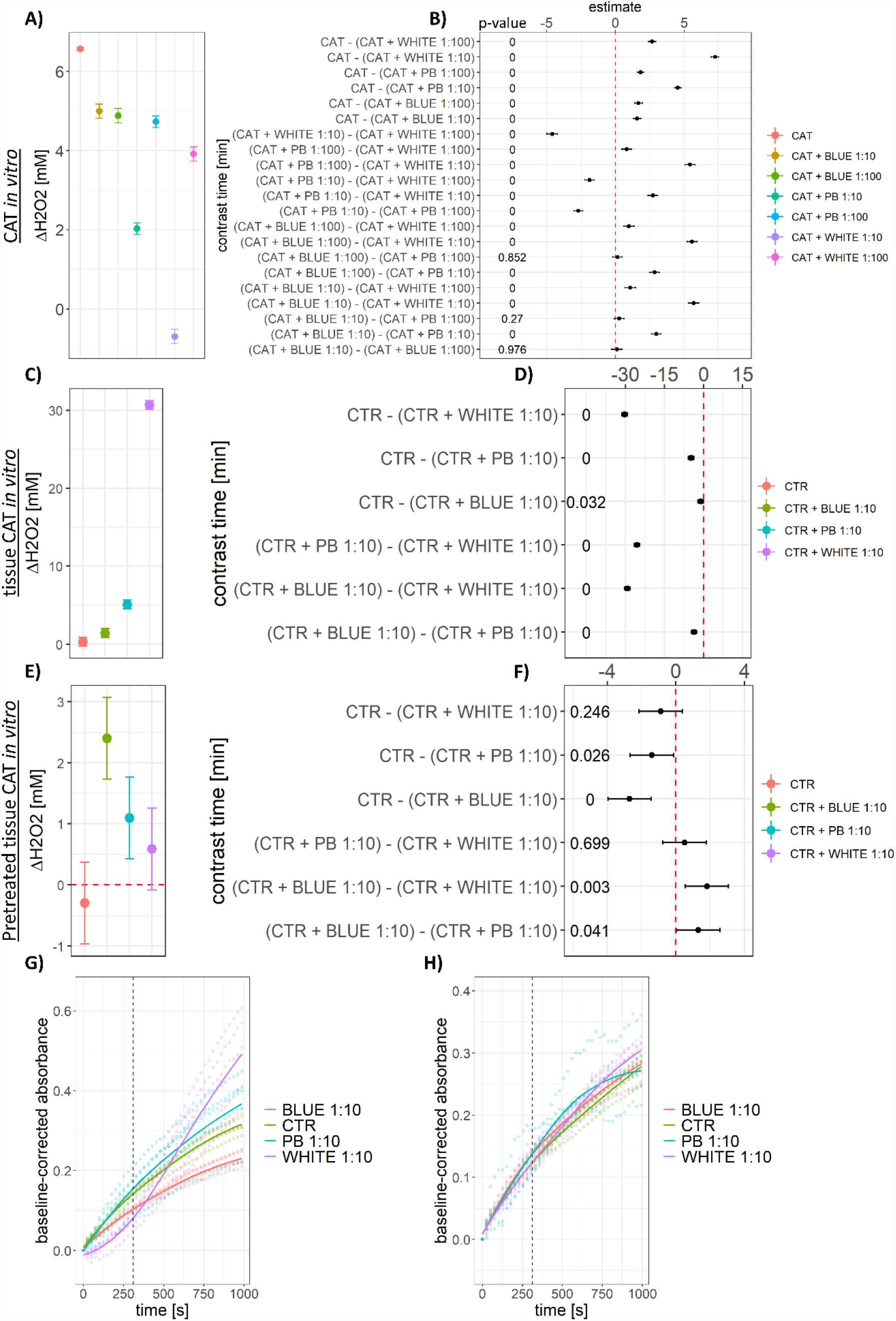
Catalase (CAT) and superoxide dismutase (SOD)-like activity of the Persian Blue tattoo ink (PB) and its blue and white fractions *in vitro* in the presence of biochemical constituents of the skin. A) Output of the model including tested dilutions of PB and its blue and white fractions and substrate concentrations (6, 8, and 10 mM H_2_O_2_), demonstrating the ability of PB and its fractions to inhibit CAT *in vitro* (t = 60 s). While the effect was not dose-dependent for the blue fraction, the white fraction and PB showed a pronounced dose-dependent inhibition of the enzyme. B) Model results presented as point estimates of differences of estimated marginal means with 95 % confidence intervals for the model presented in A. C) Output of the model including tested dilutions of PB and the blue and white PB fractions demonstrating the ability of the samples to potentiate CAT activity in the complex biochemical matrix of normal skin tissue (substrate concentration: 10 mM H_2_O_2_; t = 600 s). D) Model results presented as point estimates of differences of estimated marginal means with 95 % confidence intervals for the model presented in C. E) Output of the model including tested dilutions of PB and the blue and white PB fractions demonstrating the effect of the samples on CAT activity in the complex biochemical matrix of normal skin tissue following pre-incubation of the dyes with the tissue samples (pre-incubation time: 180 min; pre-incubation temperature: 37 °C; substrate concentration: 10 mM H_2_O_2_; t = 600 s). F) Model results presented as point estimates of differences of estimated marginal means with 95 % confidence intervals for the model presented in E. G) SOD mimetic activity of PB and the blue and white PB fractions in the complex biochemical matrix of normal skin tissue. At t < 300 s, PB demonstrated no SOD-like activity, while both blue and white fractions acted as SOD mimetics. At t > 300 s, PB was associated with slight potentiation, while the white PB fraction induced pronounced autooxidation of 1,2,3-trihydroxybenzene. Conversely, the blue PB fraction demonstrated SOD-mimetic activity. H) SOD mimetic activity of PB and the blue and white PB fractions in the complex biochemical matrix of normal skin tissue following incubation of the dyes with skin tissue for 180 min at 37 °C. Pre-incubation of the dyes with the skin tissue constituents alleviated the effects observed in F.

## Discussion

The presented results support the hypothesis that PB, a blue CuPC-based tattoo ink, can act as a mimetic of CAT and SOD and provide a possible mechanistic explanation for the reduced levels of oxidative stress in the skin with a blue tattoo [9]. Although CuPC can act as a dual functional mimetic of CAT and SOD and suppress lipid peroxidation *in vitro* [17], it was hypothesized that the *a priori* assumption that PB would necessarily reflect the properties of its main component (CuPC) would be unjustified considering the unspecified concentration of CuPC and the presence of other chemicals that may theoretically annihilate or even reverse its potential antioxidant effects. The results indicate the caution was reasonable. Although PB demonstrated CAT and SOD-like activity, the observed H_2_O_2_ dissociation capacity was relatively modest, and SOD-like activity was present only in the first part of the assay and demonstrated high variability across trials (**Fig 1**). Furthermore, the CAT-mimetic action was not persuasive once PB was fractionated (**Fig 2C-E**), while the SOD-like activity of the fractions (**Fig 2F**) suggested that large variability and time-dependence observed in the first experiment (**Fig 1B**) may have reflected the opposing effects of chemical constituents on THB autoxidation. Despite the initial hypothesis that CuPC may be the main chemical constituent of tattoo ink responsible for the effects of a blue tattoo on skin redox homeostasis [9], potentiation of THB autoxidation by the white PB fraction (**Fig 2F**) indicated that there are likely at least two chemical mediators with possibly opposing actions. Although it was not possible to confirm the presence of individual chemical constituents in PB fractions, it is highly likely that BaSO_4_ and TiO_2_ were the main constituents of the white fraction, and that they may be responsible for the observed potentiation of THB autoxidation. Although both BaSO_4_ and TiO_2_ can induce oxidative stress in different models [27,28], TiO_2_ may be a more likely mediator of the observed effect, as it stimulates the expression of antioxidant defense systems to a greater extent *in vitro* [29]. In addition, TiO_2_ can generate superoxide radicals and other ROS by reducing oxygen, due to an increased number of conduction band electrons following light exposure [26,30]. In the context of the effects of blue and white PB fractions on SOD activity, the reported time-dependence of SOD-mimetic properties of PB (**Fig 1B**) may be related to the limited ability of blue fraction constituents to suppress THB autoxidation, potentiated by the chemicals present in the white fraction of the ink.

In addition to the observed CAT/SOD-mimetic activity *in vitro*, in order to affect redox balance *in vivo*, tattoo ink should be able to exert the effect in the presence of the complex biochemical matrix of normal skin tissue (i.e in the presence of endogenous CAT, SOD, and regulators of their activity). In the presence of CAT, PB inhibited the H_2_O_2_ dissociation rate with the most pronounced inhibition observed with the white ink fraction (**Fig 3A, B**). Although little is known about the effects of BaSO_4_ on CAT activity, it has been shown that TiO_2_ can bind to CAT via electrostatic and hydrogen bonding forces, destabilize its structure and affect its activity in a dose-dependent manner [31]. Interestingly, the effects of tattoo ink on CAT were drastically altered in the presence of the biochemical constituents of normal skin, as all tested samples potentiated the relatively low endogenous H_2_O_2_ dissociation potential (**Fig 3C, D**). The white fraction of the ink induced the most pronounced effect, increasing the activity 78-fold, while the blue fraction induced only "a modest” ∼5-fold increment. Interestingly, the potentiation of CAT observed with the unfractionated ink was somewhere in between (∼17-fold), indicating that chemical constituents of tattoo ink might either act as competitive activators or engage in some other form of interaction that is reflected in CAT activity. A similar pattern was observed regarding the SOD-mimetic action, where the constituents from the blue and white fractions exhibited SOD-like properties on their own but canceled each other out when added to the tissue homogenate together (PB) (**Fig 3G**). Why were both ink fractions able to potentiate SOD activity, and whether the observed effect was mediated by the intrinsic SOD-mimetic properties of the chemical constituents or their action on the endogenous enzyme, defies a simple explanation and remains to be further explored. Nevertheless, considering that TiO_2_ can potentiate the activity of SOD [32], one possibility is that TiO_2_ may exert a dose-dependent modulatory effect on SOD similarly as has been shown for CAT [31].

Finally, the effects of PB and its fractions have been tested upon prolonged (180 min) incubation with skin homogenates at homeothermic temperature (37 °C), to assess whether some of the effects may be transient (e.g. due to dependence on an endogenous substrate) and affected by physiological temperature. Interestingly, both CAT and SOD-mimetic properties were dramatically altered by pre-incubation and the most pronounced H_2_O_2_ dissociation rate was observed for the blue ink fraction followed by PB (**Fig 3E, F**). The SOD-mimicking effect was annihilated by the pre-incubation procedure (**Fig 3H**). The exact nature of the observed phenomenon and whether the homeothermic pre-incubation more faithfully reflects the fate of the tattoo ink constituents in the human skin remains to be elucidated. On one hand, prolonged incubation at physiological temperature may provide more accurate environmental conditions for the biochemical reactions that may take place in the human body. On the other hand, the observed potentiation of the enzymatic activity may be dependent on a particular substrate present in the biochemical matrix of the tissue homogenate in limited quantities (in contrast to its continuous influx *in vivo*). Another possible explanation for the observed discrepancy might be a temperature-induced change in physicochemical properties of the tattoo ink constituents. Tattoo inks contain nanoparticles and both CuPC and TiO_2_ can be found in the nanoparticle form in the blue tattoo ink (with the mean diameter of the CuPC/TiO_2_ being 167 nm for the Intenze™ blue tattoo ink)[33,34]. Nanoparticles have an intrinsic potential to generate ROS, which has been recognized as a key mediator of nanotoxicity [35]. Considering that toxicity, photoreactivity, and ROS-generating potential depend on the particle size, shape, surface characteristics, and the crystal structure [30], a hypothetical reaction between the tattoo ink and either the tissue homogenate or the microtiter plate that may alter the structure of its nanoparticle components may provide an explanation for the observed alteration of the modulatory activity on CAT and SOD (**Fig 3**).

In the context of previously reported redox-related changes in the skin with a blue tattoo, the results presented here support the hypothesis that the suppression of oxidative stress in the N-of-1 study may be related to the antioxidant properties of some constituents present in blue tattoo ink [9]. In the N-of-1 study, a 15% reduction in lipid peroxidation in the blue tattoo was associated with 11.8% greater CAT activity [9]. In concordance with this, in this study, the constituents of blue tattoo ink were able to potentiate CAT activity in the presence of the biochemical components of the skin (**Fig 3C-F**). Interestingly, in the same study, SOD activity was slightly higher in the tattoo sample; however, this result was taken as highly uncertain considering the difference was in the range of the coefficient of variation of the method [9]. In the context of the *in vitro* results presented here, it can be assumed that apart from the limited precision of the utilized method, SOD activity in the blue tattoo may have been unchanged due to the opposing action of different constituents of the tattoo ink (**Fig 3G**) or due to the same phenomenon responsible for the loss of SOD-mimetic activity following prolonged incubation at 37°C (**Fig 3H**).

## Conclusion

The presented results confirm the hypothesis that blue, CuPC-based tattoo ink can act as a CAT and SOD mimetic *in vitro* and provide evidence that the antioxidant effects of a blue tattoo *in vivo* [9] may be mediated by the ability of CuPC and other chemical constituents of the blue tattoo ink to mimic the activity of endogenous antioxidant enzymes. In contrast to the assumption that the CAT and SOD-mimetic properties of the tattoo ink would be primarily explained by the presence of CuPC, the results suggest that both the CuPC-enriched blue and the residual white fraction may exert CAT and SOD-mimetic properties and affect redox balance, indicating that other chemical constituents (e.g. TiO_2_) may also be involved in modulation of redox homeostasis. Finally, it has been demonstrated that the ability of different constituents of the tattoo ink to potentiate and/or inhibit the activity of CAT and SOD depends on numerous factors (e.g. the presence of other constituents that exert synergistic, additive, or antagonistic effects; the concentration of the constituents and/or the substrate; the presence of compounds from the biochemical matrix of the skin; the incubation time and temperature) that should be taken in account.

## Limitations

Several important limitations should be emphasized. First, it was not possible to analyze the presence, or the quantity of individual chemical constituents of the tattoo ink used in the experiment, and the presence of different chemicals was assumed based on the official material safety data sheet of the product. Apart from the CuPC that was most likely present based on the UV-Vis spectrum characterized by the Soret peak (B-band) and the Q-band with the Davydov splitting characteristic for the phthalocyanine derivatives, the existence of other chemicals (and possibly also impurities) could not have been confirmed experimentally. Furthermore, the presence of CuPC and TiO_2_ nanoparticles was assumed based on the experimental data for the Intenze™ blue tattoo ink reported by Høgsberg et al. [34], however, the existence or the size of nanoparticles was not confirmed experimentally and the potential influence of the experimental conditions on the particle aggregation, size, shape, surface characteristics, or the crystal structure (important for the redox-related effects) was not assessed. Finally, the possibility that the chemical reaction with some of the reagents may have introduced bias in some measurements can never be completely ruled out. For example, it has been observed that 450 nm absorbance of some of the dilutions of some fractions was affected by spectrophotometric measurements (possibly due to light exposure as some chemical constituents such as TiO_2_ act as well-known photocatalysts [26])(**Supplement**). Nevertheless, precautionary steps were taken to prevent the chemical bias that may have been introduced due to unforeseen chemical reactions of samples and reagents (e.g. baseline validation model was established and analyzed for each sample individually to ensure that the expected changes such as the dissociation of H_2_O_2_ can be assumed and quantified without the risk of chemical interaction) (**Supplement**).

## Funding

None.

## Conflict of interest

None.

## Data availability statement

Data can be obtained from the author’s GitHub repository (https://github.com/janhomolak).

## Author’s contributions

**JH** conceived the study, conducted the experiments, analyzed data, and wrote the manuscript.

## Notes

### Competing Interest Statement

The authors have declared no competing interest.

