## Supplement for "In vitro analysis of catalase and superoxide dismutase mimetic properties of blue tattoo ink"

Some observations pertaining to the characteristics of the Persian blue (PB) tattoo ink were made in the experiments that did not make it to the final draft of the manuscript. Considering they may provide important contextual and methodological information the observations were described here. In the first experiment, PB exerted superoxide dismutase (SOD)-like behavior at t < 300 s, but potentiated 1,2,3-trihydroxybezene (THB) autoxidation after the 300 s time-point (Fig 1B). One possible explanation for the observed effects is that some chemical constituents of PB have limited SOD-mimicking capacity that was exceeded at t > 300 s. Another explanation may be that the observation itself affected the result considering the autoxidation of THB was assessed by continuous measurement of absorbance at 450 nm using 25 light flashes/well/measurement point. As some constituents of the ink (e.g. TiO_2_) are well-known photocatalysts [1] it is possible that light exposure affected the SOD-like properties of PB. The experiment was done following the same protocol described for the estimation of SOD-like activity, however, the samples were incubated in the buffer from which the THB was omitted. Interestingly, the effect of incubation/light exposure has been observed for the 1:10 PB sample (Fig S1). Considering repeated measurements of the same samples (shown as trials) demonstrated increasing absorbance of the 1:10 PB sample upon repeated light exposure at different time points the possibility that light exposure might have introduced bias should be acknowledged and considered. A similar observation was made following PB fractionation with the most pronounced effects seen with the 1:10 dilution of the white fraction (Fig S2).

Considering the samples had different compositions both in terms of chemical constituents and their concentrations, there was a possibility that unforeseen chemical reactions between samples and reagents would be able to introduce bias and affect the obtained results. For example, if an unknown chemical from the sample reacted with the [Co(CO_3_)_3_]Co it might have introduced bias that would preclude the measurements as it would be impossible to obtain a sensible standard dilution model required for the calculation of the amount of residual H_2_O_2_ for the tested sample. For this reason, a “baseline validation model” was established for each sample by incubating the sample with the Co(NO_3_)_2_ stop solution and nominal concentrations of H_2_O_2_ at t = 0 s to obtain the model from which the concentration of H_2_O_2_ could be estimated. Examples of such models for two dilutions of the blue and white PB fractions are shown in Fig S3. Individual baseline models were used for the calculation of residual H_2_O_2_ for each sample as calculating the difference in H_2_O_2_ from the difference in absorbance would introduce a substantial bias as evident from the fact that regression coefficients (slopes) were affected by samples being tested. The latter represents an important methodological caveat that should be considered in future research of similar design.

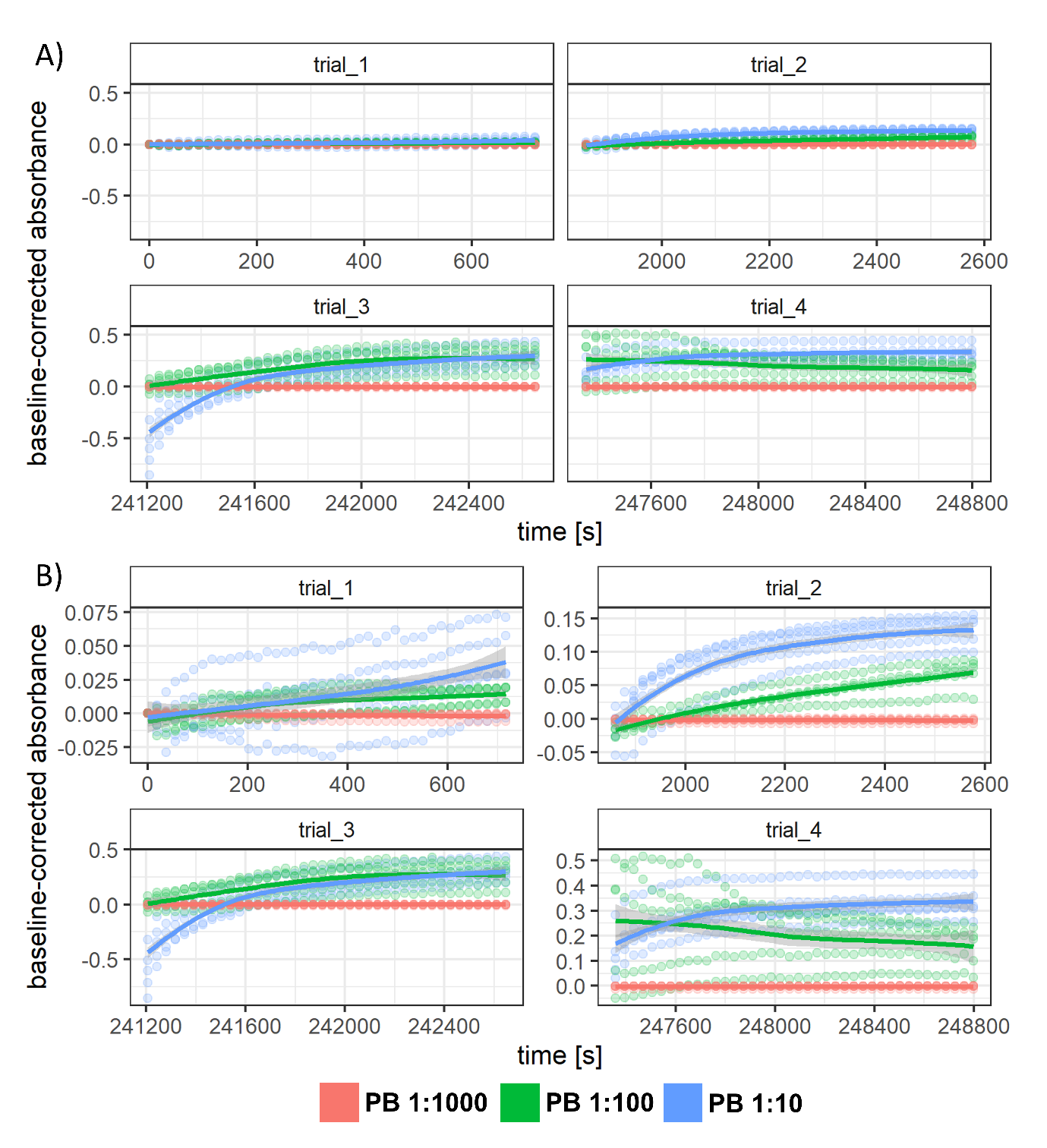

Fig S1. Baseline-corrected repeated absorbance measurements at 450 nm for different dilutions of the Persian blue (PB) tattoo ink shown with fixed (A) and free y-axes (B).

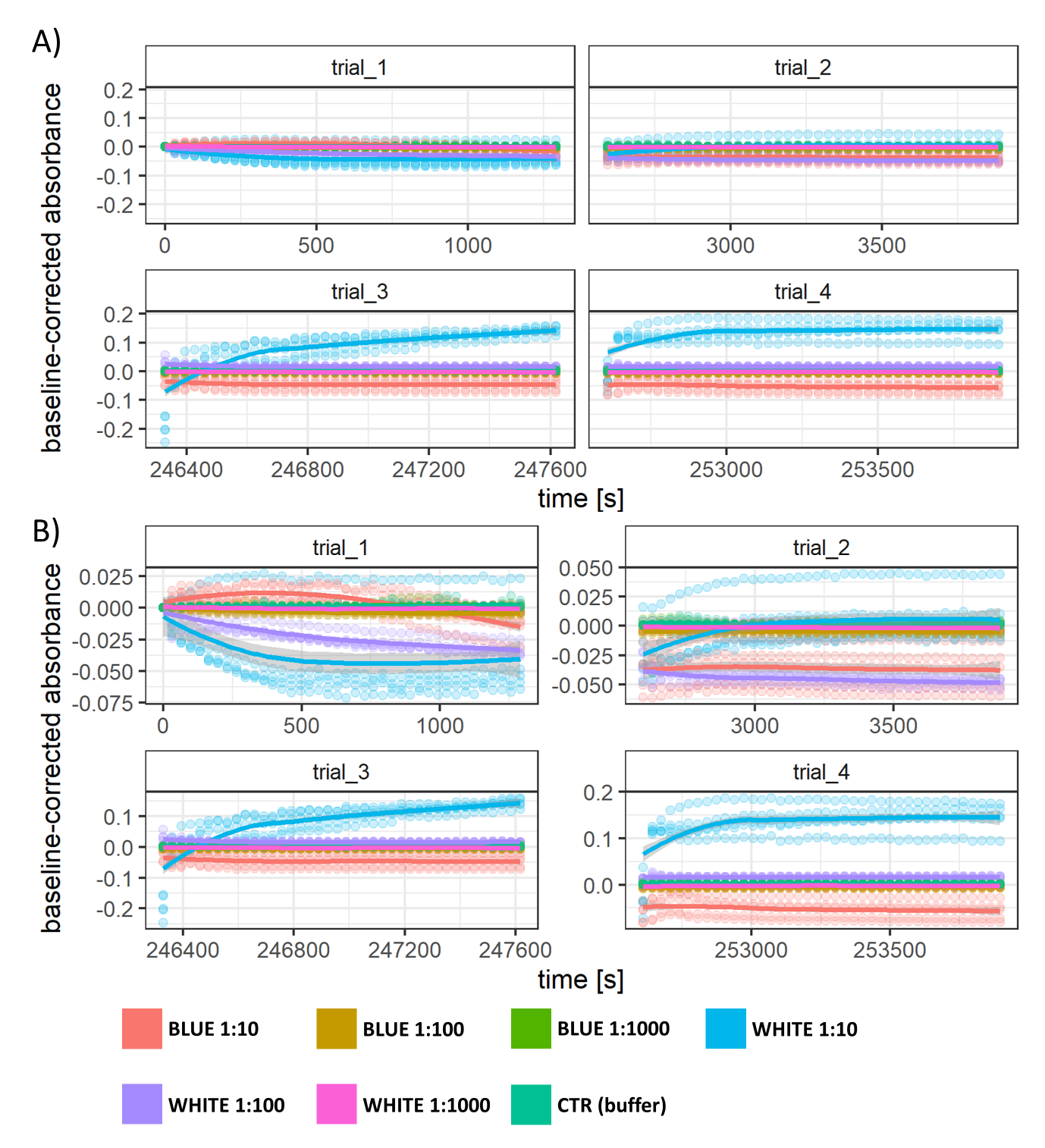

Fig S1. Baseline-corrected repeated absorbance measurements at 450 nm for different dilutions of the Persian blue tattoo ink blue and white fractions shown with fixed (A) and free y-axes (B).

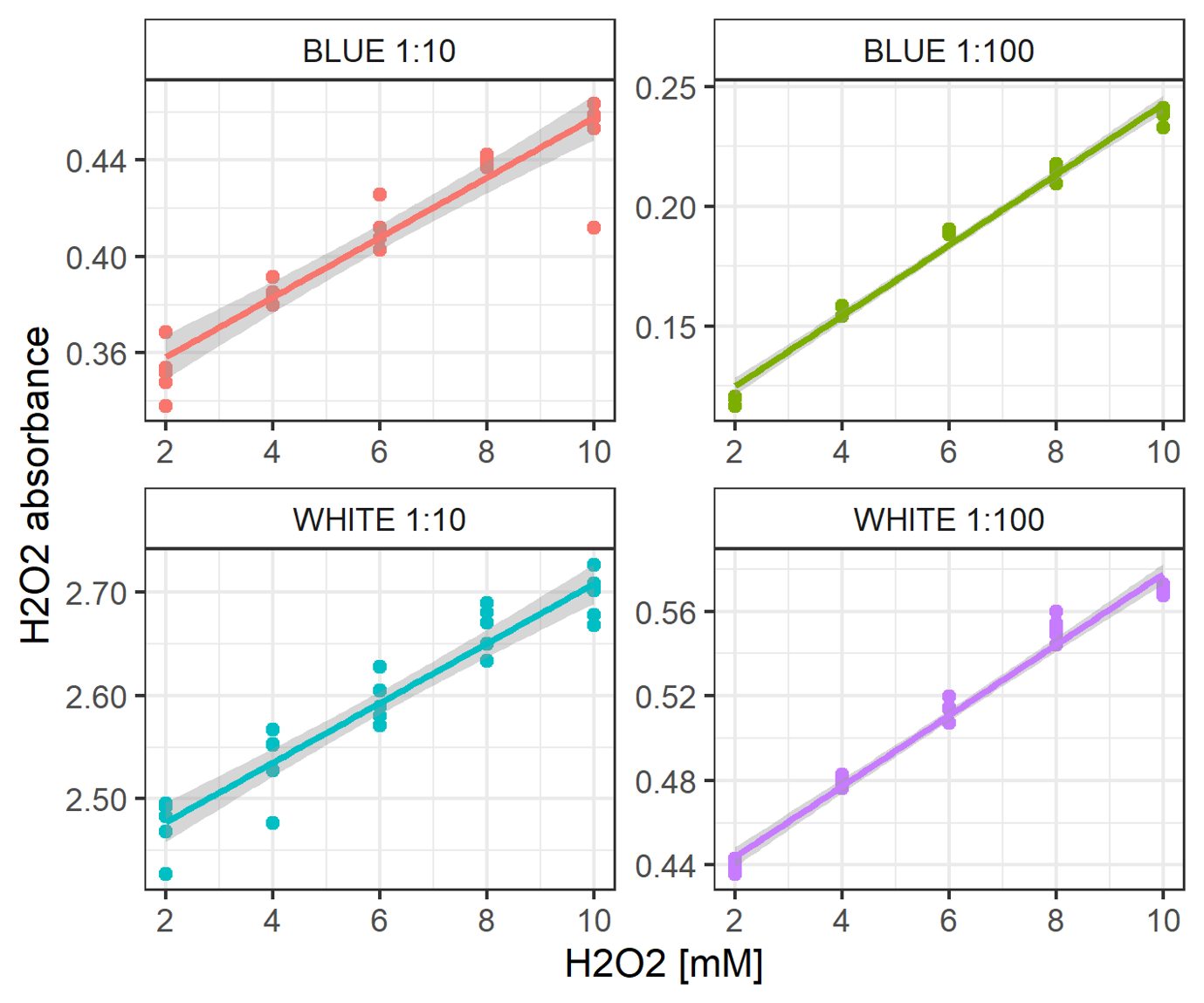

Fig S3. Baseline validation models for two different dilutions of the blue and white fraction of the Persian blue tattoo ink.
